# A Glycan-Aware Diffusion Model for Carbohydrate and Glycoprotein Structure Prediction

**DOI:** 10.64898/2026.07.16.738959

**Authors:** Keshav Sundar, Hyunjun Yang

## Abstract

Biomolecular diffusion models can now predict proteins and heterogeneous complexes, but glycans remain difficult because their branched topology, conformational flexibility, and strict stereochemical rules must be captured simultaneously. We developed SweetFold, a glycan-aware adaptation of Boltz-1x for the structure prediction of free glycans, glycoproteins, and protein–glycan complexes. SweetFold represents glycans as pseudo-polymers rather than generic ligands, preserving monosaccharide identity, anomeric state, glycosidic connectivity, and atom-level stereochemistry. We pair this representation with glycan-specific architecture, stereochemical supervision, and a sugar-centric training curriculum. Across monosaccharide, oligosaccharide, lectin, and glycoprotein benchmarks, SweetFold improves structural metrics relative to baseline all-atom diffusion models while retaining protein-only benchmark performance. These results show that chemically localized representation and supervision can extend biomolecular diffusion models to carbohydrate chemistry.

---

Glycans expand the activity landscape of proteins, lipids, and cell surfaces.^1–5^ They regulate folding, trafficking, stability, immune recognition, receptor engagement, cell–cell communication, and the behavior of biologics. Unlike proteins, glycans are not synthesized from a linear genetic template. Instead, glycoforms emerge from enzyme expression, substrate availability, organelle trafficking, and local cellular state.^6–8^ This makes the glycome biologically informative, heterogeneous, context-dependent, and difficult to categorize.^9–12^

The same features that make glycans powerful biological regulators complicate 3D glycan modeling. A glycan can consist of various combinations of monosaccharides, anomeric states, linkage positions, and branch topologies while rotating around glycosidic bonds to sample several low-energy conformations. Experimental glycan structures are often poorly resolved and partially modeled, leading to less uniformity when compared to their protein counterparts. Carbohydrate modelling tools such as GlycoShape^13^, GLYCAM-Web^14^, and CHARMM-GUI Glycan Modeler^15^ enable the sampling of glycan conformational landscapes. GlycoShape is particularly useful for training because it provides structural ensembles from MD trajectories, rather than a single static structure.^13^ Its simulations are initialized through GLYCAM-Web^15^ and parameterized with the carbohydrate-specific GLYCAM06j-1 force field^16^, providing physically plausible torsion distributions for flexible glycans.^13^. However, high-throughput prediction of glycan chemistry, protein–glycan interfaces, and glycoprotein structure remains a major challenge.^17,18^

Biomolecular models have elevated the baseline for structural biology. AlphaFold2^19^ and RaptorX^20^, and subsequent diffusion models including AlphaFold3^21^, Boltz-1^22^ and Boltz-2^23^, and Chai-1^24^ enable prediction of heterogeneous assemblies of proteins, nucleic acids, ligands, ions, and non-canonical amino acids. However, glycans expose a mismatch between these architectures and carbohydrate chemistry. When glycans are encoded as generic ligands, SMILES graphs^25^ or Chemical Component Dictionary (CCD) monomers^26^, they fail to robustly represent monosaccharide identity, anomeric state (α/β configuration), glycosylation topology (*N*-, *O*-), and stereochemistry. This gap is due to a lack of glycan-centric tokenization, architecture, and training curriculum.

SweetFold is a glycan-aware adaptation of Boltz-1x^22^ that treats glycans as first class molecules. SweetFold enables structure prediction of oligosaccharides, lectins, and glycoproteins while providing a general strategy for adapting biomolecular models to molecular classes where current representations are insufficient to meet complex chemical needs.

## Results

### Glycan Tokenization

We begin SweetFold by tokenizing glycans as pseudo-polymers. Rather than treating glycans as generic ligands, which typically use SMILES notation, SweetFold parses condensed glycan IUPAC notation^27^ into annotated glycan chains. These chains consist of atomically-tokenized monosaccharides that retain residue identity, anomeric state, branching, and glycosidic connectivity (**Fig. 1**). This pseudo-polymer tokenization scheme enables the model to recognize recurring monosaccharide residues across different contexts while preserving the atom-level chemical resolution required for accurate structure prediction. In contrast, SMILES-based inputs represent glycans as a large single residue object, leading to context dilution in the atom attention encoder.

**Figure 1.**
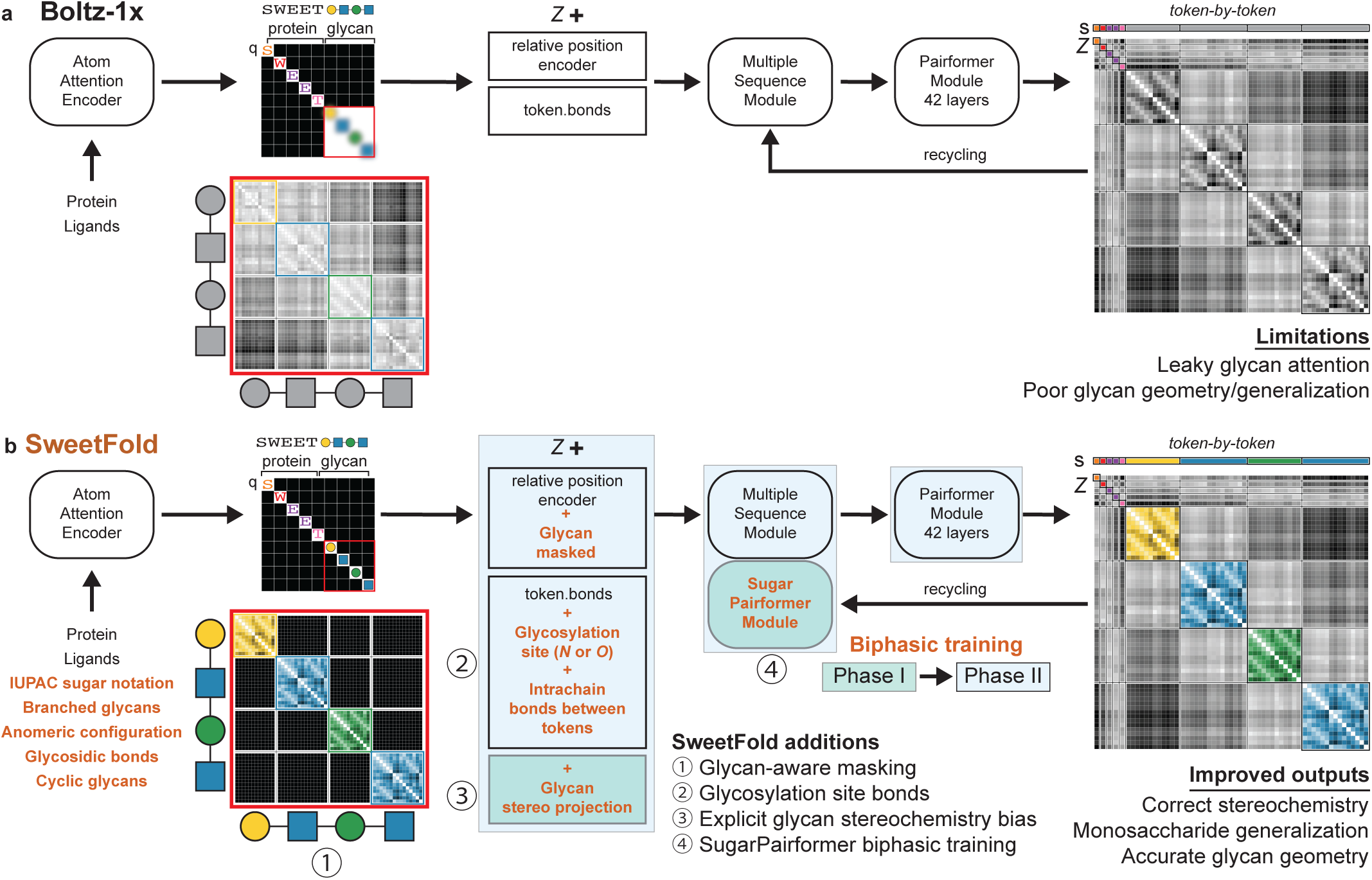
Glycan-aware tokenization and architecture in SweetFold. (a) In Boltz-1x, glycans are represented as generic ligands and enter the Atom Attention Encoder without residue-level glycan partitions, leading to context dilution and poor generalization. (b) SweetFold represents glycans as pseudo-polymers parsed from IUPAC glycan notations, preserving monosaccharide identity, anomeric state, glycosidic bonds, cyclic glycans, and glycosylation site bonds. SweetFold introduces four glycan-specific modifications: (1) Glycan-aware masking, (2) explicit glycosylation site bonds, (3) a glycan stereochemistry projection bias, and (4) SugarPairformer enabled biphasic training. These additions mask monosaccharide attention, encode local stereochemistry in the pair representation, and enable sugar-centric learning while retaining the core Boltz-1x architecture. Together, these modifications improve stereochemical validity, monosaccharide generalization, and glycan geometry.

### Architectural Modifications

To explicitly encode glycan stereochemistry in Boltz-1x, SweetFold adds a monosaccharide-focused multi-layer perceptron (MLP)^28^ called the stereo_projection bias, alongside the token.bonds bias. Monosaccharide-specific features and atom name information are projected into the *Z*-representation of sugars. This provides the model with information needed to distinguish closely related monosaccharides, including epimers and anomers.

After the modification of the single and pairwise representations, the SugarPairformer updates sugar-specific representations. The SugarPairformer module was introduced to support the biphasic training schedule and to provide context-agnostic weights for learning glycan chemistry regardless of whether the glycan is free-floating, non-covalently bound to a lectin, or covalently attached to a protein.

#### Training Strategy

Biphasic training was critical for improving training efficiency. Phase I consisted of free glycans and CCD monosaccharides, and all model weights except those involved with the stereo_projection bias and SugarPairformer were frozen. This stage allowed the model to learn monosaccharide stereochemistry and glycosidic geometry while retaining the protein folding ability inherited from Boltz-1x marked by CASP15 control benchmark (**Figure S3**). In Phase II, the full training distribution was introduced, consisting of free floating glycans, monosaccharides, lectin–glycan complexes, and glycoproteins.

To further support prediction of rare glycan residues, a custom training sampler was used. The sampler cropped and balanced *N*-linked glycoproteins, *O*-linked serine glycoproteins, *O*-linked threonine glycoproteins, lectin complexes, free glycans, and monosaccharides during Phase II training. The sampler enabled the model to learn rare structural features such as *O*-linked glycosylation.

In addition to sampling, we introduced three glycan-focused auxiliary losses to guide structure prediction; a Linkage loss for glycosidic and protein–glycan bonds; the Glyco_AA_MSE loss to prevent distortion of glycosylated amino acids; and a Dihedral loss to improve glycan substituent and glycosidic torsion accuracy.

#### Data Handling

SweetFold was trained using PDB structures^29^, GlycoShape MD-trajectory clustered oligosaccharide conformations^13^, and CCD monosaccharide structures. (**Data S1 and S2**). GlycoShape data was likely important for teaching SweetFold the conformational landscape of glycans, something that PDB data alone cannot provide.

To meet SweetFold’s additional data requirements, a new preprocessing pipeline was created. The pipeline treats glycans as pseudo-polymers while extracting monosaccharide-level CCD identity, glycosidic linkage patterns, protein–glycan attachment sites, and anomeric states. The pipeline also corrects common data errors, including inconsistent CCD assignments and residue name mismatches, to maximize data availability and stereochemical consistency (**Data S1-S7**).

### Free-floating Monosaccharide Prediction

To evaluate SweetFold’s ability to learn local carbohydrate chemistry, we constructed a free-floating monosaccharide benchmark. For all AlphaFold3 (AF3) predictions, the BAP (BondedAtomPair) input format was used (**Data S8**)^18^. The monosaccharide benchmark (**Fig. 2**) highlights the improvement of SweetFold over AF3 and Boltz-1x for free-floating monosaccharide CCD codes present in the PDB. Performance was evaluated across three metrics: Sugar-RMSD < 0.25 Å, which reports the fraction of models meeting a structural accuracy threshold; Dihedral Error < 0.1 rad, which measures the fraction of structures whose substituent dihedrals fall within 0.1 radians of the ground truth; and Total Glycan Stereochemistry, which captures the fraction of samples with perfect stereochemical accuracy. Boltz-1x was not assigned Total Glycan Stereochemistry score as glycans are treated as one large residue, thus preventing ground truth stereochemistry assignments. SweetFold outperformed both Boltz-1x and AF3 across all three-scoring metrics.

**Figure 2.**
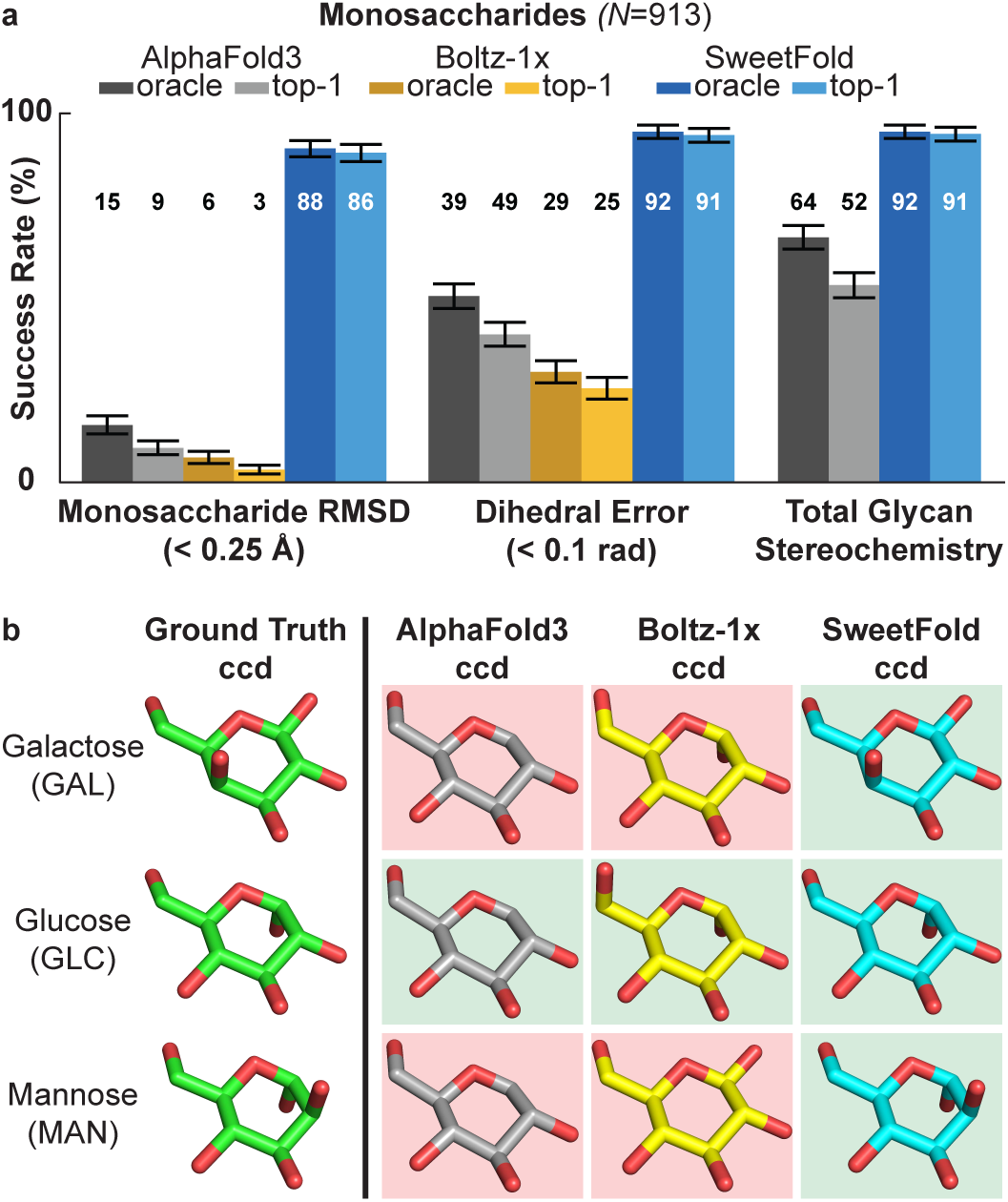
SweetFold improves monosaccharide prediction. (a) Benchmark of scored glycan CCD monosaccharides (*N*=913) using Sugar-RMSD, substituent dihedral error, and total glycan stereochemical validity. AF3 predictions used BAP notation. For each metric, oracle selects the prediction with the best metric score and top-1 selects the highest confidence score. Error bars indicate 95% confidence interval computed by bootstrap resampling over benchmark targets. (b) Representative predictions for galactose, glucose, and mannose show that SweetFold preserves sugar-specific stereochemistry, whereas AF3 and Boltz-1x collapse distinct monosaccharides into glucose-like outputs, likely because it is the most common monosaccharide in available structural data.

Representative predictions for galactose, glucose, and mannose illustrate the failure mode of generic representations (**Fig. 2b**). AF3 and Boltz-1x tended to collapse galactose and mannose and other CCD monosaccharides toward glucose, whereas SweetFold captured the stereochemical differences that distinguish these monosaccharides. This improvement is consistent with the explicit encoding of monosaccharide identity and stereochemistry through pseudo-polymer tokenization and stereo_projection. Without monosaccharide type and atom name information before the Pairformer, sugars with similar compositions but different stereochemistry can collapse into most trained common entities. Training on individual CCD codes also enabled over-sampling of rare monosaccharides, further explaining SweetFold’s ability to accurately predict free-floating monosaccharides.

### Oligosaccharide Benchmarks

Oligosaccharides extend the monosaccharide problem by adding branching, anomeric diversity, and glycosidic torsional flexibility. Small changes in stereochemistry or linkage geometry can substantially alter glycan structure, and many biologically relevant glycans sample several low-energy conformations. Experimentally resolved complex oligosaccharides are rarely found in the PDB and can include weakly resolved residues. For these reasons, global RMSD alone can be noisy or difficult to interpret. We therefore emphasized stereochemical validity and torsion-aware structural comparisons for oligosaccharide benchmarking.

We benchmarked oligosaccharide prediction capabilities on 34 structures (**Data S9**) (**Fig. 3a**). Generalization was assessed using *de novo* glycans not found in training data (**Data S10**). SweetFold generated a higher fraction of stereochemically valid structures than AF3 on both benchmarks. The larger improvement on *de novo* glycans indicates that SweetFold learned transferable sugar-level information rather than only memorizing complete glycan sequences present in training. CCD monosaccharides provided local stereochemical definitions, whereas GlycoShape MD conformers provided plausible oligosaccharide shapes and glycosidic torsion distributions.

**Figure 3.**
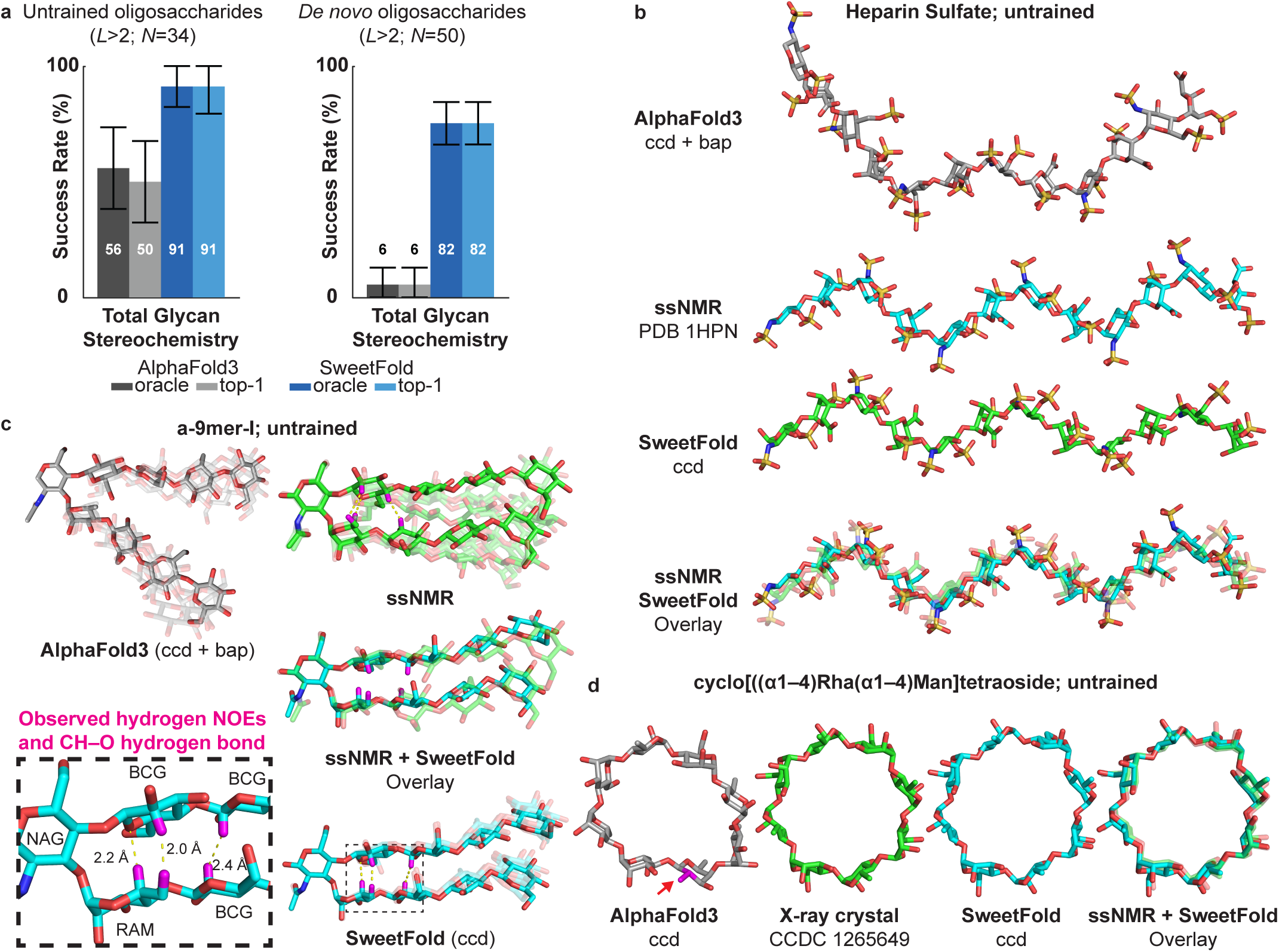
SweetFold predicts stereochemically plausible oligosaccharides preserving stereochemistry, glycosidic torsions, and hydrogen bonding patterns. (a) Total glycan stereochemistry on untrained and *de novo* oligosaccharide benchmarks. SweetFold substantially improves stereochemical validity compared to AF3, with the largest gain observed for *de novo* glycans. Error bars indicate 95% confidence interval computed by bootstrap resampling over benchmark targets. (b) Prediction of an untrained heparin sulfate compared with an ssNMR-supported structure (PDB 1HPN). SweetFold recovers an extended conformation and hydrogen bonding pattern closer to the experimental model; [–4]IDS(α1–4)SGN(α–)_6._^30^ (c) Prediction of an untrained glycan hairpin a-9mer-I compared with a ssNMR-supported structure. SweetFold reproduces the compact hairpin-like geometry and the observed sugar–sugar hydrogen bonding interaction; BGC(b1-4)BGC(b1-4)BGC(b1-4)RAM(a1-3)[BGC(b1-4)BGC(b1-4)BGC(b1-4)BGC(b1-4)]NAG(b).^31^ (d) Prediction of a synthetic cyclic rhamnose–mannose oligosaccharide compared with the crystallographic structure. SweetFold preserves cyclic topology, glycosidic geometry, and stereochemistry, whereas AF3 produces distorted stereochemistry and linkage geometry. Stereochemical error in AF3 is highlighted in magenta with a red arrow.; RAM(α1– 4){MAN(α1–4)RAM(α1–4)MAN(α1–4)RAM(α1–4)MAN(α1–4)RAM(α1–4)MAN(α1–4)RAM(α1–4)MAN(α1–4)}.^32^

Structural examples show that these gains extend beyond local stereochemistry. For a heparan sulfate sequence with solution-state NMR (ssNMR) supported ground truth^30^, SweetFold recovered an extended conformation and hydrogen bonding network resembling the experimental model, whereas AF3 lost this network (**Fig. 3b**). For a glycan hairpin stabilized by specific sugar substitutions^31^, SweetFold recovered the hydrogen bond between rhamnose and glucose that closes the hairpin and matched the ssNMR-supported MD ensemble more closely than AF3 (**Fig. 3c**). SweetFold also recovered the torsion angles and stereochemistry of a synthetic cyclic glycan consisting of rhamnose and mannose repeats^32^, whereas AF3 generated distorted stereochemistry and linkage geometry (**Fig. 3d**).

These results suggest that the CCD and GlycoShape structures play complementary roles. CCD data defines the structure and stereochemistry of monosaccharides, while GlycoShape MD structures expose the model to conformational ensembles of flexible oligosaccharides. Because GlycoShape conformers are derived from clustered MD trajectories parameterized with GLYCAM06j-1 carbohydrate parameters and MD clusters, they are useful for teaching plausible glycosidic torsions and sugar-ring arrangements.^13,14,16^ This combination was particularly important for *de novo* glycans, which have no complete experimental ground truth counterpart.

### Lectin–Glycan Benchmarks

Lectins provide a difficult test for glycan-aware structure prediction because protein structure, glycan structure, and non-covalent protein–glycan interactions must be modeled together. Lectin–glycan data is comparatively scarce, and glycan density in experimental structures is often incomplete or weakly resolved. A useful benchmark therefore needs to score both protein structure and glycan-centric properties rather than relying only on global protein accuracy and local ligand docking.

We evaluated lectin structure prediction using two benchmarks—a PDB benchmark of resolved lectin–glycan complexes (**Data S11**) and an untrained benchmark assembled from UNIPROT lectin sequences paired with biologically relevant oligosaccharides (**Data S12**). On the PDB benchmark, SweetFold and AF3 achieved similar global protein accuracy and glycan scores, whereas Boltz-1x performed worse, consistent with the limitations of generic SMILES-based glycan input for protein–glycan complexes (**Fig. 4a**).

**Figure 4.**
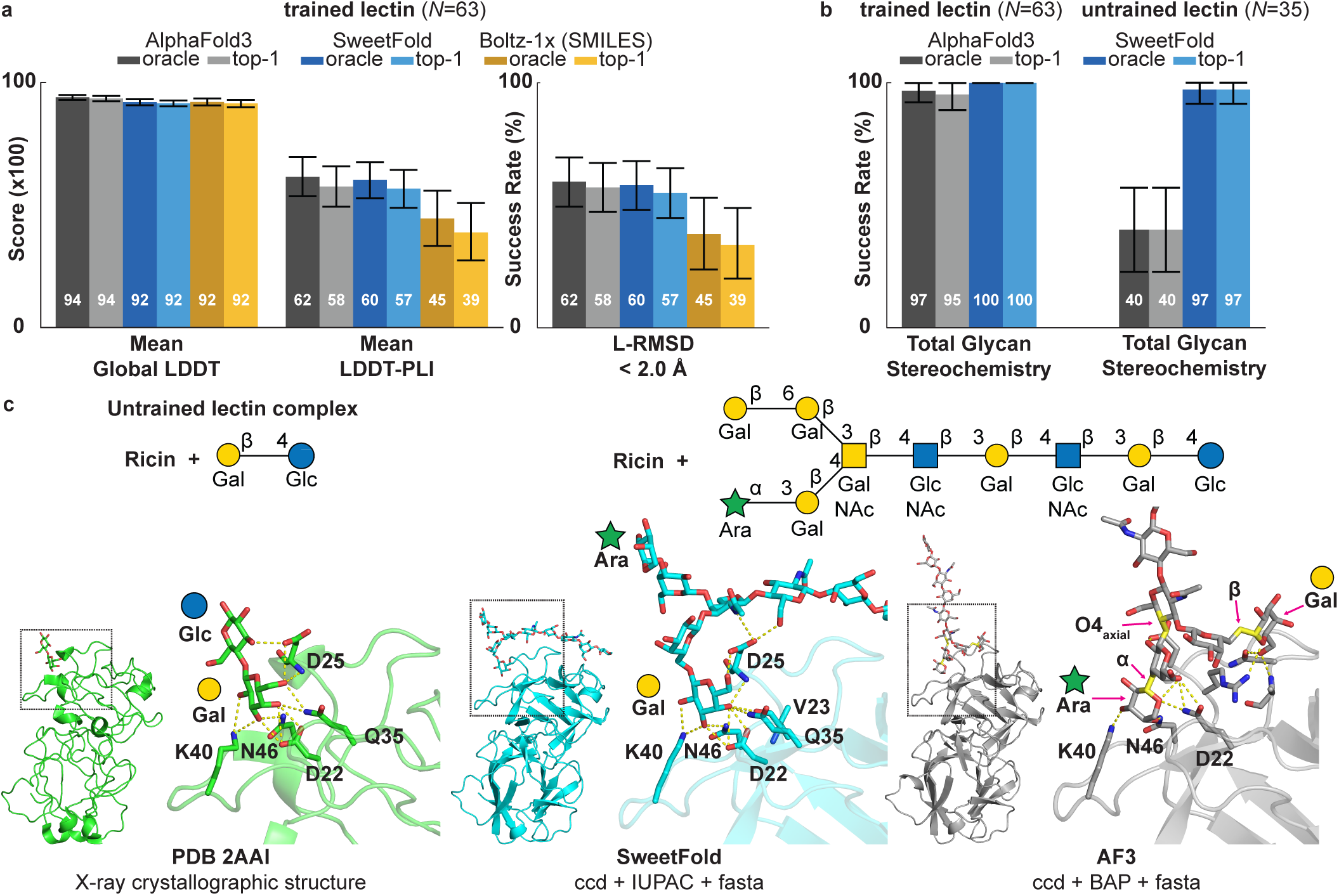
SweetFold preserves glycan stereochemistry in lectin–glycan complex prediction. (a) Performance on PDB lectin–glycan complexes comparing global protein plDDT, protein–ligand interaction scores, and ligand RMSD success rate. SweetFold maintains protein accuracy comparable to AF3 while improving over Boltz-1x. Error bars indicate 95% confidence interval computed by bootstrap resampling over benchmark targets. (b) Total glycan stereochemistry success rates for trained and untrained lectin–glycan benchmarks. SweetFold preserves glycan stereochemistry across both benchmarks, including untrained lectin-associated glycans. (c) Representative untrained ricin–glycan prediction. The crystallographic ricin–disaccharide complex shows recognition of terminal galactose in the binding pocket (PDB 2AAI). SweetFold places the terminal galactose (Gal) in a compatible binding pose, whereas AF3 places an incorrect arabinose (Ara) residue with multiple stereochemical errors in the pocket shown in yellow with red arrows. This example illustrates how local glycan stereochemistry can influence protein–glycan recognition.

Because experimental complex structures were not available for the untrained lectin benchmark, we scored glycan stereochemical plausibility rather than ligand placement. SweetFold retained higher stereochemical validity when compared to AF3 predictions (**Fig. 4b**). This result suggests that pseudo-polymer tokenization, stereo_projection bias, and SugarPairformer condition the diffusion trajectory that generalizes to unseen lectin-associated glycan.

Ricin was used as an example to illustrate how glycan stereochemistry can affect protein-glycan interactions (**Fig. 4c**). Ricin is a crystallographically characterized lectin that recognizes terminal galactose residues. In the PDB file 2AAI, ricin is crystalized with a BGC–GAL disaccharide.^33^ When the ricin protein sequence was paired with an untrained glycan sequence, SweetFold placed the terminal galactose in the expected binding pocket. In contrast, AF3 placed a stereochemically incorrect arabinose (ARA) residue in the pocket.

These examples suggest that local glycan stereochemistry can be coupled to protein– glycan recognition. Errors in anomeric state or substituent orientation can change which residues interact with a lectin surface. By preserving monosaccharide identity, anomeric configuration, and glycosidic geometry, SweetFold generates more plausible lectin–glycan poses, particularly with regards to larger, untrained oligosaccharides.

### Glycoprotein Benchmark

Glycoproteins extend the glycan prediction problem from non-covalent recognition to covalent protein–glycan chemistry. Accurate prediction requires the model to correctly position the root monosaccharide, preserve stereochemical rules, and avoid distortion of the glycosylated amino acid. SweetFold therefore augments glycosylation site learning through Glyco_AA_MSE loss while also introducing protein–glycan inter-chain covalent bonds.

We evaluated glycoprotein prediction on a PDB glycoprotein benchmark (**Data S13**) and two untrained *N*-and *O*-glycosylation glycoprotein benchmarks (**Data S14**). The untrained benchmarks used UNIPROT derived *N*-and *O*-glycoprotein sequences with biologically relevant glycosylation sites and glycan sequences not observed in the training dataset. In addition to global lDDT, ligand RMSD, protein–ligand interaction score, and total glycan stereochemistry (**Fig. 5a)**, we evaluated glycoprotein specific geometry using the Privateer glycosite torsion score^34,35^. For Privateer scores, we report the fraction of samples with no potential torsion outliers using a threshold of −1.0 for each evaluated site.^35^ Glycosylation site geometry was also validated using angle-based scores (**Fig. 5b**). For *N*-glycosylated glycans, the ASN CG–ASN ND2–glycan C1 angle was required to fall between 110° and 130°.^2^ For *O*-glycosylated glycans, Ser CB–Ser OG– Glycan C1 or THR CB–THR OG1–glycan C1 angle was required to fall between 104° and 124°.^2^

**Figure 5.**
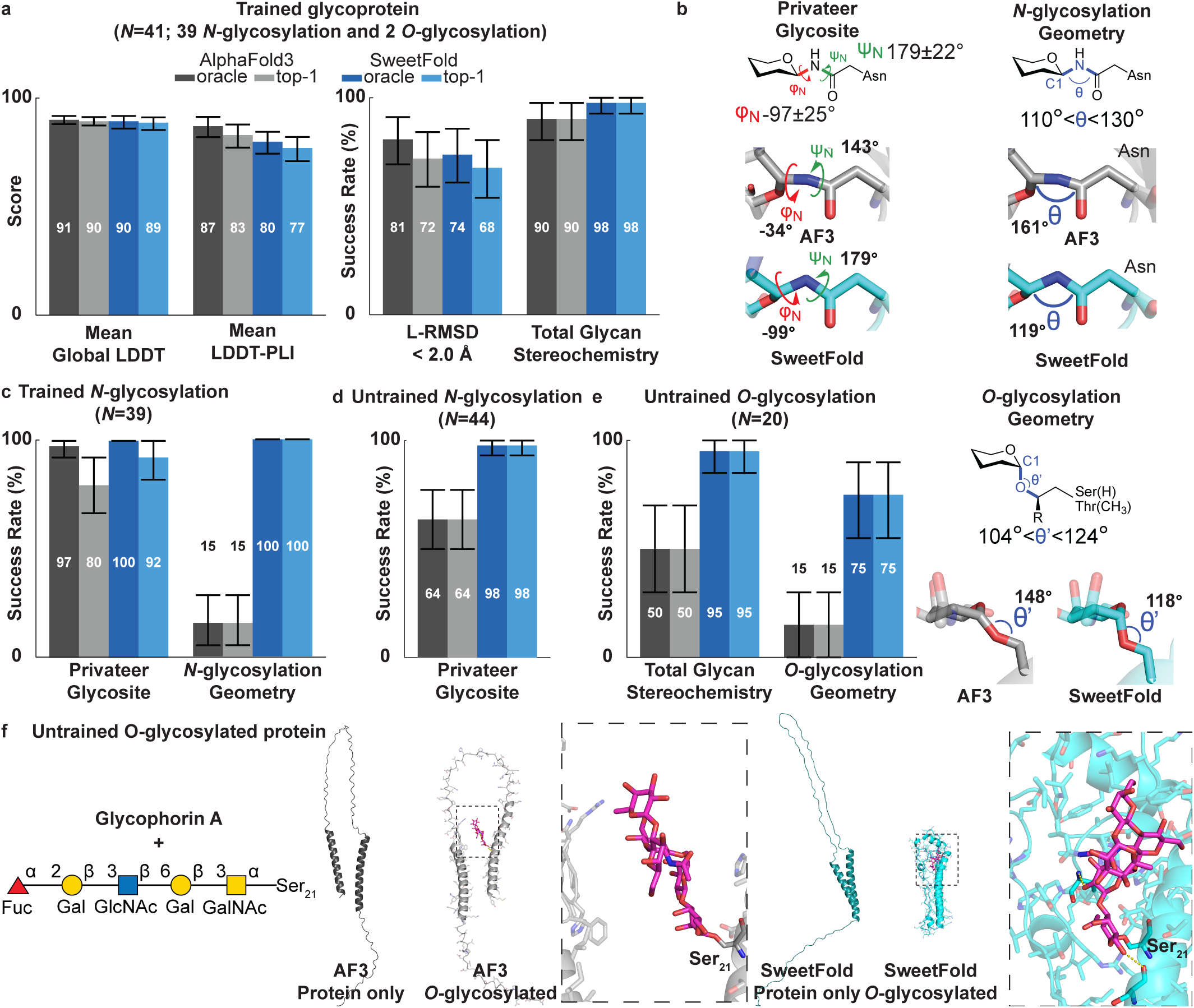
SweetFold improves glycoprotein stereochemistry and glycosylation site geometry. (a) PDB glycoprotein benchmark (*N*=41) comparing global lDDT, protein–ligand interaction score, ligand RMSD, and total glycan stereochemistry. Both models retain strong global protein accuracy, but SweetFold improves glycan centric validity. Error bars indicate 95% confidence interval computed by bootstrap resampling over benchmark targets. (b) Glycosylation site geometry metrics used to evaluate *N*-glycoprotein predictions. Privateer glycosite torsions assess the local N-glycan linkage conformation, and *N*-glycosylation geometry evaluates whether the Asn-linked glycosidic angle falls within the expected range. Representative examples show that SweetFold preserves glycosite torsions and Asn-linkage geometry more accurately than AF3. (c) Trained *N*-glycosylation benchmark showing improved Privateer glycosite validity and *N*-glycosylation geometry for SweetFold. (d) Untrained *N*-glycosylation benchmark showing that SweetFold retains high glycosite torsion validity on glycoproteins not observed during training. (e) untrained *O*-glycosylation benchmark comparing total glycan stereochemistry and *O*-glycosylation geometry. SweetFold improves both root glycan stereochemistry and Ser/Thr-linked glycosidic geometry relative to AF3. (f) Representative untrained *O*-glycosylated glycophorin A prediction. SweetFold predicts glycan-dependent local organization surrounding glycosylated residue Ser21, whereas AF3 shows less glycan-dependent structural prediction. Together, these results indicate that SweetFold improves glycan stereochemistry, glycosite torsion accuracy, and *N*-and *O*-glycosylation site geometry while retaining strong global protein structure performance.

On the PDB glycoprotein benchmark, both AF3 and SweetFold performed well on global structure scores and glycan stereochemistry (**Fig. 5a**). However, glycosylation site geometry metrics highlight a major difference between the models. AF3 frequently distorts the glycosidic bond angles at the glycosylated amino, resulting in poor geometric validity (**Fig. 5b,c**). SweetFold preserved the expected ASN-C1 geometry, consistent with the Glyco_AA_MSE loss.

The difference was more apparent on untrained glycoproteins. On the untrained *N*-glycoprotein benchmark, SweetFold maintained high Privateer glycosite validity and *N*-glycosylation geometry, whereas AF3 showed a large drop in glycosite plausibility (**Fig. 5d**). On the untrained *O*-glycosylation protein benchmark, SweetFold retained high glycan stereochemical accuracy as well as generating more plausible *O*-glycosylation geometry compared to AF3 (**Fig. 5e**). This improvement is consistent with the glycan centered training curriculum, which oversampled rare *O*-glycosylated structures during training.

SweetFold also captured glycan-dependent local protein changes in an untrained example (**Fig. 5f**). In the glycosylated protein prediction of glycophorin A, a nearby disordered protein segment folded toward the glycan and formed polar contacts—in the protein only prediction, the same region remained disordered. AF3 predicted a similar protein conformation with and without the glycan. Although this example does not have an experimental ground truth structure, it illustrates how improved glycan stereochemistry and linkage geometry can influence local protein organization around a glycosylation site.

Together, these benchmarks show that accurate glycoprotein prediction requires more than placing a ligand near a protein surface. The model must learn the covalent linkage, root sugar stereochemistry, glycosite torsion, and local amino acid geometry simultaneously. SweetFold improved these features across *N*-and *O*-glycosylated proteins while retaining strong protein structure folding performance (**Figure S3**).

## Discussion

SweetFold indicates that the main barrier to glycan prediction in biomolecular diffusion models is not data scarcity alone, but also how carbohydrate chemistry is represented and learned. Although glycans are sparse in structural databases, this scarcity becomes severely limiting when they are treated as diffuse collections of atoms or generic ligand graphs. SweetFold treats glycans as first class pseudo-polymers, preserving monosaccharide identity, anomeric state, branching, glycosidic connectivity and atom-level stereochemistry throughout the model. This representation allows the model to learn recurring sugar units across free floating glycans, lectin–glycan complexes, and glycoproteins while retaining the local chemical information needed to distinguish epimers, anomers, linkage positions, and substituent orientation.

The results also identify architectural features required for accurate carbohydrate structure prediction. In a generic all-atom diffusion framework, the features and token.bonds bias alone are too weak to define complex glycan geometry, especially when identical atomic composition can correspond to different epimers or anomers. The stereo_projection bias addresses this limitation by adding sugar specific atom pair priors that encode monosaccharide identity and stereochemical context before the diffusion. The SugarPairformer then updates sugar-level pair features in a context-agnostic manner, allowing glycan chemistry learned from free-floating monosaccharides and oligosaccharides to transfer to lectin complexed and protein-linked glycans. Together, these components make it possible to train glycan specific parameters without disrupting the inherited protein folding capability of the base model.

The training curriculum was equally important. Because sugars are rare relative to proteins in the PDB, a standard training distribution would rapidly dilute glycan containing examples. SweetFold therefore uses a biphasic curriculum where phase I focuses on monosaccharide and free-floating glycans, while phase II introduces the full distribution of free-floating glycans, lectins, and *N*-and *O*-linked glycoproteins. Glycan-centered cropping and sampling further prevent common protein rich examples from overwhelming rare carbohydrate classes. These design choices suggest that efficient learning of rare biomolecular chemistries requires sampling and supervision at the chemical unit of interest, rather than solely at the complex level.

The benchmark results support this interpretation across increasing levels of structural complexity. SweetFold improved monosaccharide prediction by preserving total glycan stereochemistry and substituent torsions, and the benefits extended to oligosaccharides where branching, glycosidic torsions, and low energy conformational ensembles become important. On lectin benchmarks, SweetFold maintained glycan stereochemical validity in trained and untrained complexes, and the ricin example illustrates how local sugar stereochemistry can affect which residue is presented to a protein binding pocket. On glycoprotein benchmarks, SweetFold improved glycosite torsion, root sugar geometry, and *N*-and *O*-glycosylation site validity while retaining strong protein structure prediction.

The data sources used for SweetFold appear to play complementary roles. CCD monosaccharide structures anchor the model’s understanding of individual sugar residues. GlycoShape MD conformers provide plausible oligosaccharide shapes and glycosidic torsion distributions that are difficult to infer from sparse crystallographic examples alone. This combination is especially useful for *de novo* glycans, where no identical training structure exists and the model must generalize from local stereochemical rules and physically plausible oligosaccharide conformations.

Several limitations remain. Heavily glycosylated proteins are still challenging because dense glycans introduce large conformational search spaces and increase the likelihood of steric clashes between neighboring glycans or between glycans and the contacting protein residues. Protein–sugar interactions also remain difficult for all current models because many lectin and glycoprotein contacts are partially resolved experimentally attributed to weak, transient interactions. Ground truth structures impose an additional limitation as glycans are often weakly resolved and often modeled as one member of a conformational ensemble. These issues make it difficult to distinguish model error from uncertainty in the experimental reference, particularly for flexible terminal residues and crowded glycosylation sites.

Future work should increase the amount and diversity of glycan-aware training data. One direction is to seed training with MD trajectories of lectin–glycan complexes and glycoproteins, rather than only free oligosaccharides, so that the model can learn plausible protein–sugar contacts and binding site geometry. A second direction is to use SweetFold predictions to prioritize *de novo* glycans for experimental structure validation. These improvements should also enable *de novo* lectin design, in which new protein scaffolds are designed to recognize specific glycan epitopes. Finally, the strategy developed here should extend beyond glycans. Other stereochemically complex ligands and pseudo-polymers may benefit from the same combination of chemically correct tokenization, specific pair representations architecture, and training curricula that keep rare chemical classes visible during training. In this sense, SweetFold provides not only a model for carbohydrate structure prediction, but also a route for adapting biomolecular diffusion models to rare molecular classes that are poorly represented by generic ligand tokens.

## ONLINE CONTENT

### Data availability

The training and benchmark data are available through Zenodo repository. (https://doi.org/10.5281/zenodo.21105358).

### Code availability

Source code, preprocessing scripts, training configuration files, and inference utilities are available through the SweetFold GitHub repository. (https://github.com/Keshav-Sundar-4/SweetFold/tree/main).

### Model availability

The final trained checkpoint is deposited on the SweetFold Hugging Face model page. (https://huggingface.co/Keshav-Sundar-4/SweetFold/tree/main).

A Google Colab notebook for IUPAC glycan and protein input preparation and SweetFold inference is provided here. (https://colab.research.google.com/drive/1zzUXvhls35VAOowoXYuzKW9IAYYZh1XH)

### Supplementary Information

The experimental details are available here. (PDF)

## ACKNOWLEDGMENT

H.Y. acknowledges support from the National Science Foundation (DMR-2011846) and the National Institutes of Health (R00AG084926). K.S. acknowledges support from the Summer Materials Undergraduate Research Fellowship (SMURF) Program from MRSEC at Brandeis University (DMR-2011846). The authors thank Dr. Michael Hagan for generous offer to use H100 GPUs to train SweetFold, and Dr. Yibing Wu and Dr. Rian Kormos for their thoughtful discussions.

## Author Contributions

K.S. and H.Y. designed and carried out the research; K.S. and H.Y. analyzed the data; K.S. and H.Y. wrote the paper. H.Y. supervised the study. K.S. and H.Y. read and approved the manuscript.

## Supplementary Information

### Methods

#### User Accessibility

The SweetFold model checkpoint is available on Hugging Face. (https://huggingface.co/Keshav-Sundar-4/SweetFold/tree/main).

The model, benchmark, and training code are available through the SweetFold github. (https://github.com/Keshav-Sundar-4/SweetFold/tree/main).

A Google Colab notebook is also provided for IUPAC glycan and protein input generation and for running and downloading SweetFold outputs. (https://colab.research.google.com/drive/1zzUXvhls35VAOowoXYuzKW9IAYYZh1XH).

#### Compute and Data

Four NVIDIA H100 GPUs were used to train SweetFold. Two sources of ground truth structural data were used for training—the Protein Databank (PDB)^2^ and GlycoShape^1^. PDB data consisted of X-ray crystallographic structures and cryo-EM structures that contained the allotted glycan codes (**Table S1)**. This dataset contained 21,507 training/validation PDB structures and 2,128 test PDB structures (**Table S2**), including glycoprotein, lectin, and free-floating glycan data. The structure cutoff data was March 8, 2026 for PDB database (**Data S1**). To supplement experimentally resolved structures, structures from GlycoShape was used (**Data S2**)^1^. GlycoShape contains 860 oligosaccharide structures, each with multiple molecular dynamics trajectory clusters—in total, 4,323 GlycoShape structure files were used. In addition, 930 Chemical Component Dictionary (CCD) monosaccharide structures were incorporated for the model training. Phase I dataset contained 5,253 structure files (**Data S3**), and the Phase II dataset contained 26,670 structure files (**Data S4**).

#### Training Methods

##### Glycan-Centric Data Loading

SweetFold was implemented as a fork of Boltz-1x^3^ in which the training pipeline was reworked to enable accurate glycan structure prediction while retaining protein training ability. The central methodological change was to treat glycans as pseudo-polymer ligands. The data structure definitions (`type.py; training.py; const.py`), tokenizer (`tokenize/boltz.py)`, featurizer (`featurizer.py`), and conditioning modules (`model.py; confidence.py; loss/diffusion.py`) were updated to represent glycan-containing structures as pseudo-polymers with explicit atoms, residues, chains, covalent linkages, monosaccharide identities, anomeric annotations, and glycosylation site metadata.

The Boltz-1x structure datatype (`type.py`) was extended with glycan-specific categories for glycosylation sites, per-monosaccharide features, and atom-to-monosaccharide assignments in addition to the standard atom, bond, residue, chain, and bond arrays. The data loading was modified to preserve these detailed glycan annotations.

SweetFold tokenizes glycans with a pseudo-polymer strategy that preserves the multi-residue nature of glycans while retaining atomic-level resolution. In the Boltz-1x tokenizer, glycan residues were treated as non-standard and were therefore tokenized at atom-level tokenization. For these glycan residues, each atom was assigned an independent token with its own atom index, center index, distogram index, coordinates, resolution mask, and token-to-atom mapping enabling glycan residue grouping and understanding.

At the token level, SweetFold retained standard features, including token index, residue index, asymmetric chain ID, entity ID, symmetry ID, molecule type, residue type, distogram center, resolved mask, distogram mask, cyclic period, and token–token bond features. Additional glycan-specific token features were added, including token-monosaccharide type, a one-hot anomeric state, a monosaccharide mask, and an inter-glycan mask (`featurizer.py`). The anomeric state and monosaccharide type features define the 3D coordinate and the stereochemical identity of each monosaccharide and determines R/S configurations for stereocenters within a given glycan. At the atom level, the featurizer preserved atomic coordinates, atom names, elements, reference coordinates, atom masks, atom-to-token mappings, and frame indices.

##### Data Preprocessing

Structure files were first cleaned to remove all non-glycan/non-protein entities (`phase1_cleaner.py`). This filter handled PDB files containing multiple structural states and examples containing glycans with clashing atoms. Atom clashes were defined as atom pairs separated by less than 1.0 Å. Glycans containing such clashes were removed to prevent training instabilities. A subsequent preprocessing script (`preprocess_glycans.py`) converted raw PDB files into the format required by the training pipeline. This preprocessing step served as the main bridge between structural files and the model’s internal feature representation. The pipeline accepted a directory of PDB files and a pickled CCD file, processed each structure, and wrote two outputs for each successfully parsed structure—(1) a compressed structure file containing atom, residue, chain, bond, connection, and atom-to-monosaccharide arrays, and (2) a JSON record describing the processed chains for inclusion in the training manifest.

GlycoShape data occasionally contained residue name mismatches, which were corrected during preprocessing. Examples of residues that often were inconsistently named in different data sources included: A2G, NAG, NGA, NDG, SIA, and TOA.

The preprocessing script first rechained filtered PDB atom records to separate glycan components to their own chains. In some structure, non-covalent glycans were grouped within the same chain, and protein-linked glycans shared a chain with their protein partner. Atom records were sorted by residue and classified as protein or glycan according to standard amino acid and CCD definitions. Glycan residues within the same chain were then linked into residue-level components.

After rechaining, the SweetFold pipeline parsed atom records into dictionaries and classified each chain as protein or glycan. Each dictionary stored the atom number, atom name, residue name, chain ID, residue number, 3D coordinates, and element. These dictionaries were used to detect both monosaccharide–monosaccharide linkages and protein–glycan glycosylation sites. The script assigned donor and acceptor directionality for detected glycosidic linkages, which were subsequently stored in the connections array.

For each detected glycosidic or protein–glycan connection, the pipeline determined the anomeric configuration of the donor monosaccharide. An atomic bond graph was generated using a distance threshold of 2.0 Å, and the main sugar ring of each monosaccharide was identified from the 3D coordinates. Within monosaccharide ring, the algorithm selected the anomeric carbon, ring oxygen, neighboring endocyclic atom, and exocyclic substituent. A dihedral angle calculated from these four atoms was then used to classify the anomeric sugar as α or β. Root and lone monosaccharides were handled separately by constructing pseudo-connections from their local anomeric substituents, which allowed the pipeline to assign anomeric information when the monosaccharide was not a child in a glycosidic bond. Dihedral angles with −95° ≤ X ≤ 95° were classified as α linkages, whereas angles outside of this range were classified as β linkages.

Two mappings—identity and anomeric mappings (`const.py; preprocess_glycans.py`)—were introduced for data correction and to maximize the SweetFold training. PDB structures occasionally contain CCD-mapping errors. For example, the CCD code *MAN*, which refers to α-mannose, is sometimes assigned as β-mannose. This error would introduce incorrect biases during the forward pass. The anomeric mapping corrected this problem by converting the base sugar and detected anomeric configuration into the corresponding α or β configuration form. A second issue is that different PDB structures assigned different CCD codes for the identical sugar, which reduces generalization across identical monosaccharide structures. The identity mapping compresses sets of identical monosaccharides into a single chemical base identity, thereby maximizing monosaccharide level data.

The data processing pipeline also applied biochemical and geometric filters to protein-linked glycans before assembling the final training structure. Candidate glycosylation sites were evaluated according to the identity of the protein residue and the detected anomeric configuration. For asparagine-linked glycans, two geometry filters were applied using atoms from the asparagine (ASN) side chain and the glycan donor atom. The script calculated the ASN CG–ASN ND2– GLYCAN C1 angle and the ASN OD1–ASN CG–ASN ND2–GLYCAN C1 dihedral angle. The candidate site was retained only when both values were within allowed thresholds (110° < X < 130° for the angle and −20° < X < 20° for the dihedral angle). ASN-linked glycans were retained only when the root monosaccharide was β-configured, whereas serine (SER)-and threonine (THR)-linked glycosylation sites were retained only when the root monosaccharide was α-configured.

#### Training Curriculum

##### Curriculum Details

SweetFold was trained under a biphasic curriculum (`train.py, full.yaml`). **Phase I** used a maximum learning rate of 0.006, while **Phase II** used a lower maximum learning rate of 0.0006. In both cases, the optimizer settings used Adam-style parameters^4^ with β1 = 0.9, β2 = 0.95, and an ε = 1e−8. The Boltz-1x^3^ training scheduler was retained to train SweetFold—a base learning rate of zero (0 step), a short warmup (5 steps), decay after a fixed number of steps, repeated decay intervals, and a multiplicative decay factor of 0.95. The larger **Phase I** learning rate was used during the initial glycan adaptation stage, in which the trainable portion of the model was restricted to the glycan-focused architecture. The lower **Phase II** learning rate was used when the broader model training distribution was introduced, including monosaccharides, free-floating glycans, lectins, and glycoproteins. **Phase I** consisted only of free-floating glycans and monosaccharides (**Table S2**) allowing the glycan-specific architecture to learn glycan biophysics and monosaccharide stereochemical rules. During **Phase I**, all model components except the *stereo_projection bias* and *SugarPairformer* were frozen. This reduced the number of trainable parameters, increased iteration speed, and minimized the risk of degrading protein folding performance by training only on sugars. A 93/3.5/3.5 training/validation/test split was used in **Phase II**. All model parameters were unfrozen in Phase II. MSAs were not used in either training phase.

**Figure S1.**
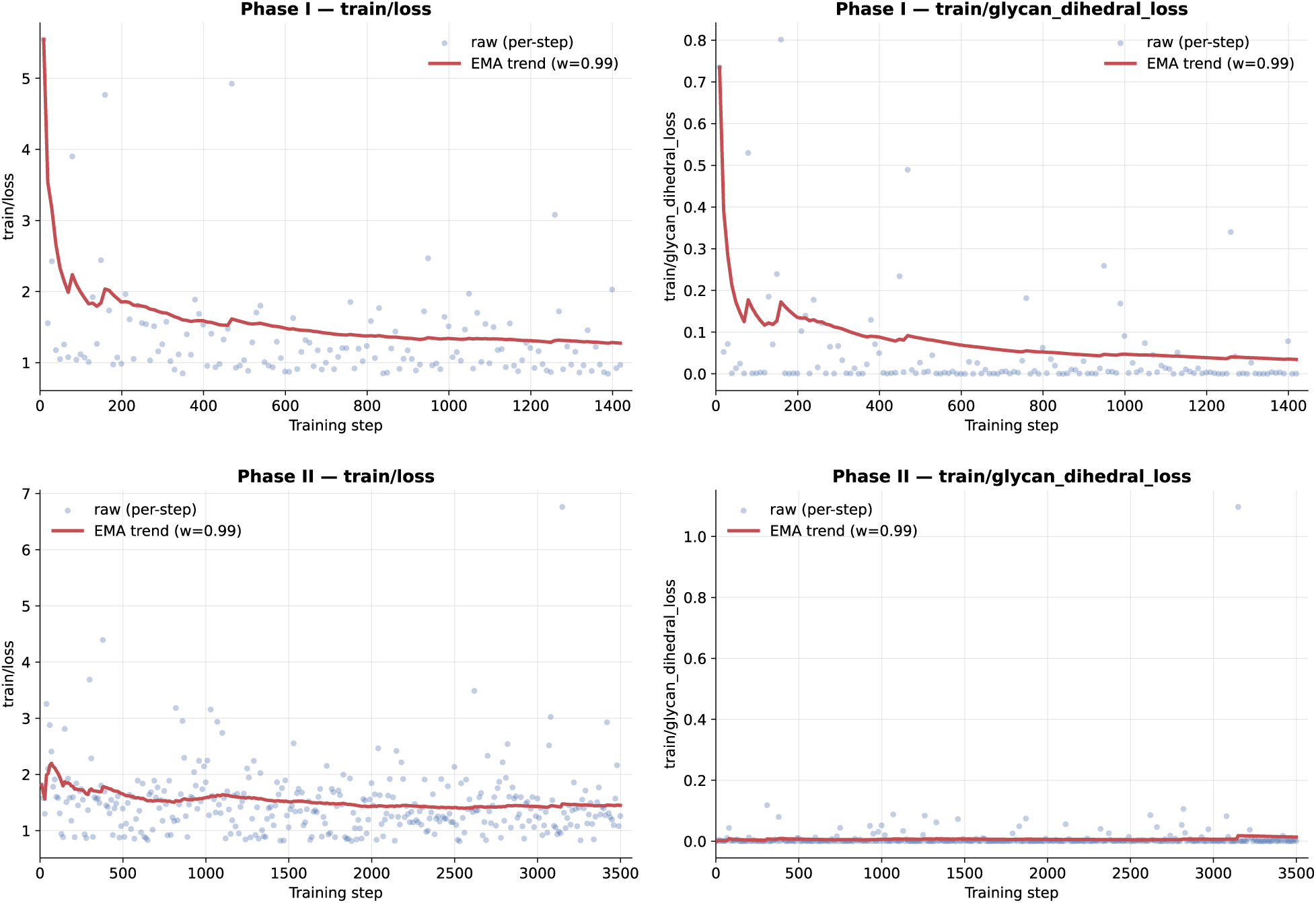
Training loss during the biphasic SweetFold training curriculum. Total loss and glycan-dihedral loss were monitored during Phase I and Phase II training. Phase I used free glycans and CCD monosaccharides to train the glycan-specific modules and showed a rapid decrease in loss during the early optimization steps, consistent with efficient learning of monosaccharide stereochemistry and glycosidic geometry. Phase II introduced the broader training distribution, including protein-containing glycan structures, and showed more leveled loss changes across the longer training schedule, likely due to the model was already heavily trained on protein structures.

##### Glycan-Centric Data Cropping

The dataset limits were chosen to support atom-level glycan modeling while controlling memory use. Training examples were capped at 320 tokens and 3,072 atoms (`full.yaml`), with atom attention operating over query windows of 32 atoms and key windows of 128 atoms. Other training hyperparameters were kept consistent with Boltz-1x. Distogram features were computed with 64 bins over distances from 2.0 Å to 22.0 Å.

Cropping (`crop/boltz.py`) was modified to center training examples on glycans rather than on random structural regions. If the complete structure fit within both the token and atom limits, the example was returned without cropping. If the structure exceeded either token or atom limits, a glycan chain was selected and a random token from that glycan chain was used as the crop center. Tokens close to this crop center in were gathered for input into training. This cropping pipeline ensured that retained crops were spatially organized around glycan content rather than around arbitrary protein region that might exclude glycans or glycosylation sites.

##### Glycan-Centric Data Sampling

A custom sampler, *equity_glycan_training* (`cluster.py`), was used for SweetFold training. Separate clusters were assigned for monosaccharides, free-floating glycans, lectins, serine-linked glycoproteins, threonine-linked glycoproteins, and asparagine-linked glycoproteins.

When equity glycan training was enabled, the sampler generated repeated chunks with fixed counts from the glycan-specific pools. Each chunk sampled 1000 monosaccharide structures, 250 free-floating glycan structures, 1000 lectin structures, 750 serine-linked glycoprotein structures, 750 threonine-linked glycoprotein structures, and 750 asparagine-linked glycoprotein structures. This was the main mechanism by which **Phase II** combined monosaccharides, free-floating glycans, lectin, glycoprotein-associated examples without requiring separate training loops for each structural class.

##### SweetFold Architectural Additions

The glycan-specific data pipeline required architectural changes so that glycan information was not only preserved in preprocessing, but also actively incorporated into the learned token and pair representations. SweetFold retained the primary Boltz1x architecture, including MSA conditioning, pair representation updates, diffusion-based structure prediction, and confidence prediction, while adding glycan-aware architectural components where needed.

##### Stereo_Projection Bias MLP (Sugar Distogram)

The glycan *stereo_projection bias* was a small multi-layer perceptron (MLP)^5^ that learned an all-to-all pair bias over glycan tokens. For each monosaccharide, the model gathers all atoms assigned to that residue and included glycosidic and glycosylated atoms connected to the donor monosaccharide. Each ordered token pair was represented using the monosaccharide type and the full atom names of both tokens. The *Stereo_Projector* mapped these features into the pair-representation dimension, producing a *K* × *K* × *token_z bias* matrix. This matrix was added directly to the global pair representation *z*, excluding self-pairs, alongside the *token.bonds bias* and *positional encoder biases* before the trunk pass. The stereochemical bias included atoms assigned to each monosaccharide and selected external glycosidic linkage atoms connected to the donor sugar. Atoms on glycosidic bonds were renamed according to the internal carbon to which they were bonded, allowing the learned bias to represent inter-residue stereochemistry.

##### Masking of the Relative Positional Encoder

The relative position encoder was modified due to the non–sequential nature of glycan topology. To mask glycan structures properly, the encoder inspected the *is-monosaccharide* feature to identify glycan tokens. For token pairs in which both tokens were glycan-associated, the *relative position bias* was suppressed by setting the projected bias to zero. This prevented glycan–glycan token pairs from receiving a sequential relative-position prior based on residue and token indices.

##### SugarPairformer

The SugarPairformer module was introduced as a refinement pathway for glycan tokens. It used the infrastructure of the standard Pairformer, with attention over both single and pairwise representations. Glycan tokens were gathered and refined exclusively, whereas protein tokens were left unchanged. This enabled for context-agnostic glycan learning and allowed the glycan specific modules to be trained independently during **Phase I** (`sugar_trunk.py`).

##### Confidence Module

The confidence module (`confidence.py`) was updated with glycan-aware components, including the *stereo_projection bias* and *SugarPairformer*. These additions allowed the confidence module to generate more accurate confidence scores for glycan structures.

##### SweetFold Training Losses

Protein–glycan glycosylation sites were converted into a symmetric glycosylation linkage mask over the *token.bond* matrix. This presentation treated protein–glycan glycosylation bonds analogous to covalent bonds within ligands and protein bonds between amino acids.

The SweetFold training objective (`loss/diffusion.py)` combined diffusion coordinate losses with additional glycan-specific geometric losses. The diffusion module remained responsible for learning to denoise atom coordinates conditioned on the trunk representations, token features, atom features, and pair features. During training, predicted coordinates were compared with aligned target coordinates using MSE loss, smooth lDDT loss, linkage loss, glycosylated amino acid MSE loss, and glycan dihedral loss.

MSE and smoothlDDT losses were retained from Boltz-1x, while three auxiliary losses were added to improve structure prediction and training efficiency. The additional loses were defined as follows;

###### Linkage loss

To enforce the correct length of the covalent protein–glycan linkage, we define;

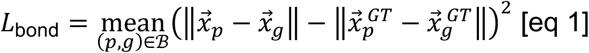

Where ℬ is the set of protein-glycan linkages, *p* is the protein atom, *g* is the glycan atom, and *GT* is the corresponding ground truth coordinates.

###### Glycosylated amino acid MSE loss

To preserve the local geometry of the modified residue and the directly attached glycan atom, we define;

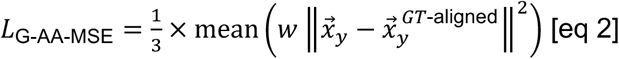

Where w is the standard *weighted_rigid_align* function from Boltz-1x and *y* is the subset of atoms consisting of the glycosylated amino acid and glycan atom (*g*) involved in a protein-glycan linkage.

###### Glycan dihedral loss

To constrain glycan ring substituent orientation, we define;

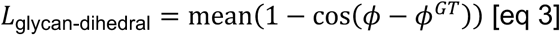

Where *ϕ* refers to the dihedral angle generated by three consecutive ring atoms and the selected substituent. For example, if the dihedral of the O2 atom in GLC was being calculated, three consecutive ring atoms (C4, C3, C2) would be gathered, and the O2 substituent. A dihedral would then be calculated using the atoms: C4-C3-C2-O2. GLC also has the following substituents atoms – O1, O3, O4, O6. The atom groups for calculating these dihedrals are shown below.

C3-C2-C1-O1 (O1)
C4-C3-C2-O2 (O2)
C5-C4-C3-O3 (O3)
O5-C5-C4-O4 (O4)
C1-O5-C5-C6 (C6)

Furthermore, for di-substituted atoms such as the C2 atom of a Sialic Acid, dihedrals are generated for both substituents. The dihedral angles for the disubstituted C2 atom of sialic acid are shown below:

C4-C3-C2-O2 and C4-C3-C2-C1

The glycosylation auxiliary loss was added to the base diffusion objective as;

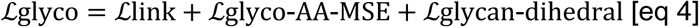

Terms with no valid sites in a batch were omitted, which is equivalent to contributing zero to the total loss.

##### Training Validation Metrics

Three glycan-focused validation metrics were included to benchmark training efficiency (`validation.py`). These included *glycan_rmsd*, *glycan_anomeric_dihedral_loss*, and *glycan_ring_dihedral_loss*. *glycan_rmsd* calculated the RMSD of the glycan portion of training example, whereas the dihedral metrics calculated the accuracy of anomeric and substituent dihedrals, respectively. The dihedral metrics used the equation described above (eq 3), except that the evaluated subsets were separated into anomeric and substituent terms to determine whether glycosidic linkages were more difficult to generate. No significant difference between the anomeric and substituent losses were observed, indicating that glycosidic linkages did not cause additional stereochemistry issues.

#### Inference Methods

SweetFold was built on top of the Boltz-1x inference path so that glycan-containing inputs could be represented as branching and cyclic structures rather than linear sequences. Proteins, DNA, and RNA can be specified as one-dimensional inputs with deterministic sequential bond topology. In contrast, glycan inputs must encode residue identity, anomeric configuration, branching, and cyclization. To accommodate these requirements, `schema.py` was modified to accept condensed IUPAC nomenclature during inference, and a parser (`schema.py`) was created to read the resulting inputs.

##### IUPAC Sequence Parsing

The IUPAC parser converts condensed IUPAC glycan notation into a molecular graph representation (`schema.py; inference.py`) that can be featurized by the SweetFold featurizer (`featurizer.py`). The parser first decomposes the input string into residues and structural annotations. Each glycan residue field contains the CCD code, anomeric configuration, and glycosidic bonds in which the residue participates. Brackets […] are interpreted as branch markers and are used to generate the corresponding glycosidic linkage notation. Parentheses (…) following a residue are treated as linkage specifications only if they contain both an anomeric configuration and numeric donor and acceptor glycan positions, such as (a1-4) or (b1-3). Root monosaccharides are represented by (a) or (b) because they do not define a donor-acceptor linkage. SweetFold applies two mappings (`const.py`), analogous to those used in the preprocessing—(1) an identity map that collapses chemically equivalent CCD codes into a defined code, and (2) an anomer map that assigns the appropriate α or β CCD form based on the input string. CIF and PDB formats require residues to be ordered sequentially, so an alternative mapping is used that flattens branching and cyclic notation into a linear sequence. This flat sequence exists solely for compatibility purposes — the branching structure itself is preserved and passed into the trunk as part of the actual feature input.

##### Condensed IUPAC Sequence Examples

**Figure S2.**
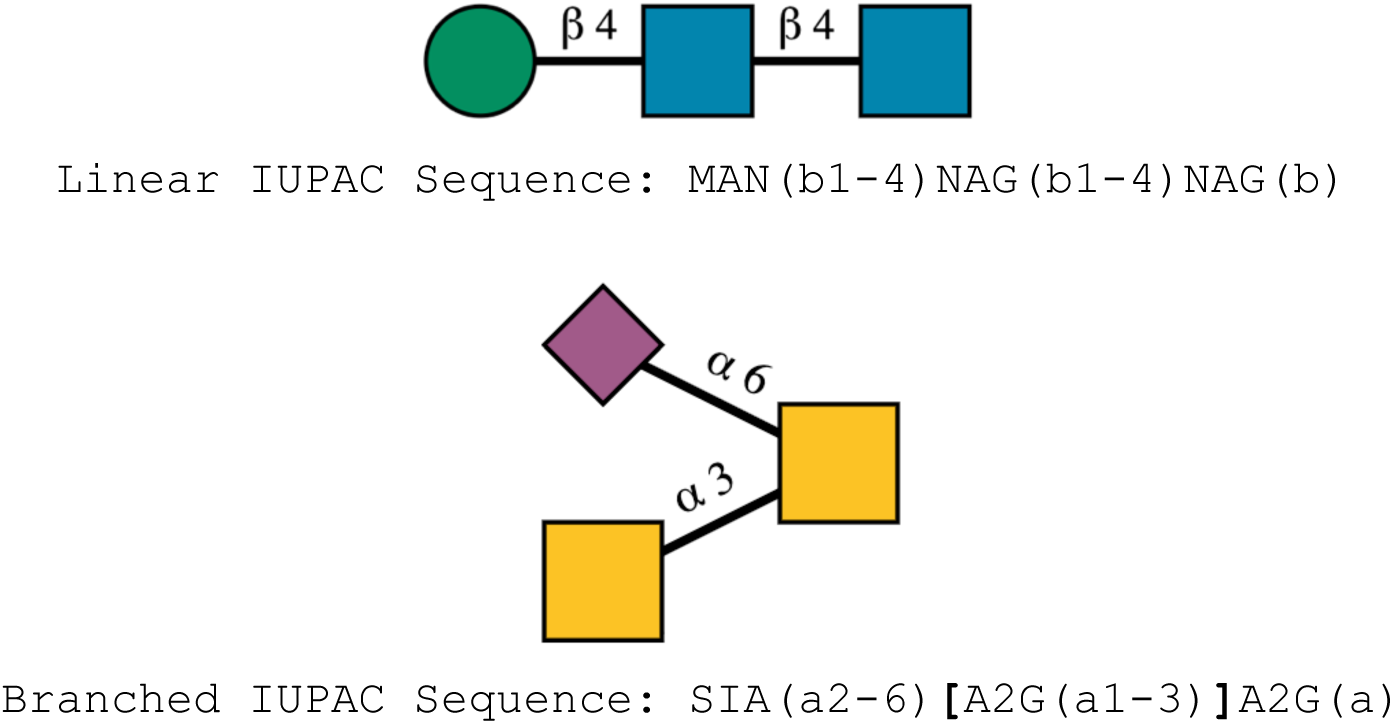
Condensed IUPAC glycan notation parsed by SweetFold. Linear and branched glycans are shown using Symbol Nomenclature for Glycans (SNFG)^6^ above the corresponding condensed IUPAC strings. Monosaccharides are identified by CCD residue codes, glycosidic linkages are written in parentheses, and branch points are represented by square brackets in the sequence.

##### Post-Sequence Parsing

After the molecular graph was generated, post-processing corrections were applied. The most important correction was the deletion of extraneous atoms created by glycosidic connections. For example, in the linkage MAN_2(_a1-4)MAN_1,_ each MAN residue initially contains an anomeric carbon (C1) and an oxygen substituent bound to that carbon (O1). After glycosidic linkage creation, C1 of MAN_2 i_s bonded to the O4 of MAN_1,_ and O1 atom of MAN_2 m_ust be removed. Post-sequence parsing performs this deletion and ensures extraneous substituents are removed from the molecule.

##### Cyclic Glycan Parsing

SweetFold also extends inference parsing to handle cyclic glycan notation. A separate parsing path was required because cyclic glycans do not contain a root monosaccharide. In the implemented notation, curly braces {…} mark the site of a cyclic bond. The parser first ignores cyclic notation and builds the glycan using the standard SweetFold parsing path. The cyclic bond is then added by connecting the two atoms forming the glycosidic bond and applying the correct anomeric configuration to the appropriate monosaccharides. Post-parsing processing steps are then applied and the final structure is passed to the featurizer.

##### Glycosylation Site Parsing

Glycosylation sites were parsed using a specialized notation that specified the chain, residue, and atom name of the connected protein and glycan components. These sites were stored in the connections array and were subsequently added to the *token.bonds* matrix.

##### Condensed IUPAC Sequence Examples

Condensed IUPAC sequence examples for cyclic IUPAC sequences and glycosylation sites. Curly brackets denote the position of a cyclic acceptor monosaccharide, as highlighted in bold. Glycosylation sites require explicit chain and residue identification, along with atom-level identification for glycan residues.

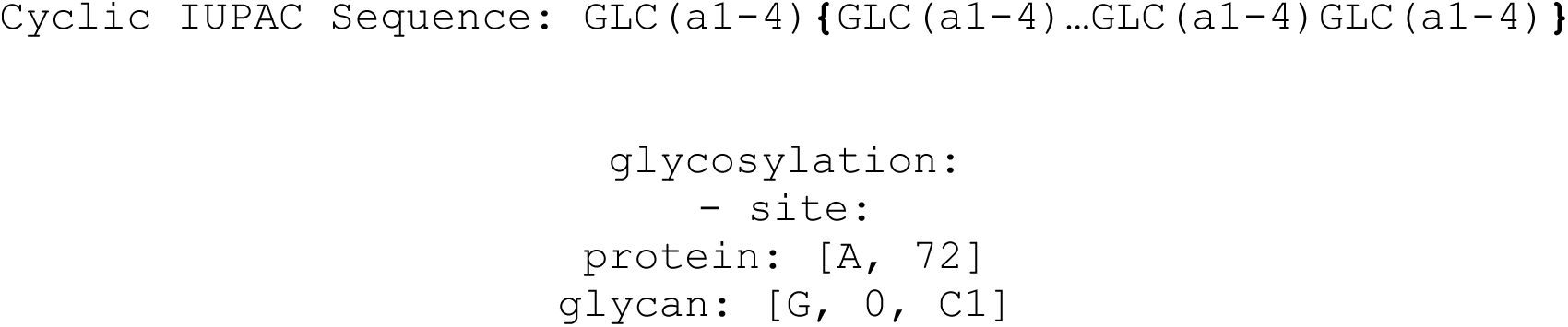

#### Benchmark Methods

For each benchmark target, inputs were generated in the format required by each prediction method. AlphaFold3 was run from BAP (Bonded Atom Pair) inputs, Boltz-1x from SMILES inputs, and SweetFold from IUPAC/CCD inputs. Each method produced predicted structures in CIF format. Evaluation scripts parsed the predicted CIF files, located the corresponding confidence files when available, and computed structure, ligand, stereochemical, and glycan-validation metrics. For reference-based protein–glycan and lectin benchmarks, global structure and ligand placement metrics were computed with OpenStructure^7^. To ensure compatibility with OpenFold temporary evaluation CIFs were created in which each glycan ligand was represented as a single non-polymer ligand component. GlobalLDDT was taken from the OpenStructure polymer comparison and reports the global local-distance difference test score between the predicted and reference macromolecular structures. LDDT-PLI^8^ was taken from the OpenStructure ligand– protein interface comparison. For each structure, LDDT-PLI values were averaged over assigned ligands. Ligand RMSD was computed after alignment using both the ligand atoms and binding pocket atoms.

##### CASP15 Benchmark

To assess whether glycan-specific training degraded protein-only structure prediction, a CASP15 control benchmark was run using SweetFold, Chai-1^9^, Boltz-1x^3^, and AlphaFold3^10^. The benchmark consisted of AlphaFold3, Chai-1, Boltz-1x, and SweetFold. SweetFold maintained protein folding performance comparable to Boltz-1x, indicating that glycan-centered curriculum did not measurably degrade the inherited protein folding capability under this benchmark.

**Figure S3.**
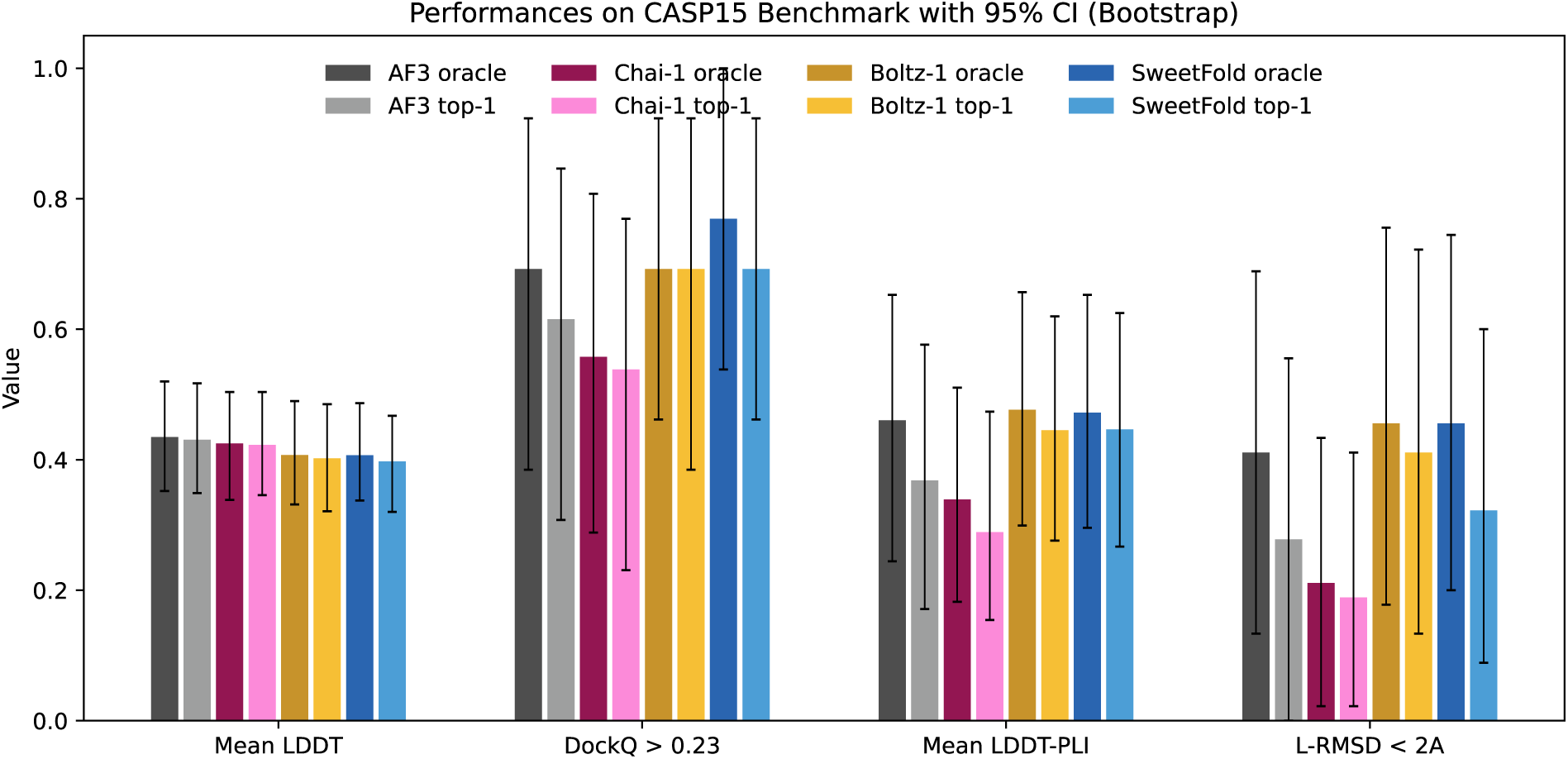
CASP15 protein only benchmark used to evaluate retention of the inherited protein structure prediction performance. AlphaFold3, Chai-1, Boltz-1x, and SweetFold were compared across LDDT, DockQC, LDDT-PLI, and L-RMSD < 2.0 Å metrics. SweetFold performed comparably to Boltz-1x, indicating that glycan-specific training did not measurably degrade protein folding, protein-ligand docking, or ligand-geometry performance in this control benchmark. Error bars indicate 95% confidence interval computed by bootstrap resampling over benchmark targets.

##### Total Glycan Stereochemistry as a Metric

Total Glycan Stereochemistry was evaluated with RDKit^11^ using monosaccharide CCD as chemistry reference. For each predicted CIF files, glycan residues were extracted and read and matched to CCD definitions by residue name. RDKit molecules were constructed from the CCD topology using the predicted heavy atom coordinates. R/S stereochemistry was assigned from the 3D coordinates and compared with the stereochemistry assigned to the corresponding CCD reference molecule. Only named R and S stereocenters that were assigned in both the predicted structure and the reference were compared. Structures with missing or unassignable stereocenters were skipped rather than counted as incorrect. A predicted sample was counted as stereochemically correct when a sample contained zero chiral errors. Total Glycan Stereochemistry is reported as the fraction of evaluated samples with zero chiral errors. This metric was computed for AlphaFold3 and SweetFold outputs; Boltz-1x was not assigned a Total Glycan Stereochemistry score because glycans are treated as one large residue, preventing ground truth stereochemistry assignment from residue level comparisons.

##### Monosaccharide RMSD as a Metric

For monosaccharide benchmarks, ligand geometry was additionally evaluated by heavy atom RMSD and substituent dihedral angle error. Heavy atom RMSD was computed after atom name matching between the predicted and reference monosaccharide and Kabsch alignment of the matched heavy atoms. Substituent dihedral errors were computed by identifying ring substituent dihedrals in the reference structure and measuring the mean absolute angle error between the corresponding predicted and reference dihedrals.

##### N-Glycosylation Site Geometric Validity

For *N*-linked glycoproteins, the ASN CG–ASN ND2–GLYCAN C1 angle was calculated at detected *N*-glycosylation sites. A structure passed the *N*-glycosylation geometry check when all detected and scorable *N*-glycosylated ASN sites had ASN CG–ASN ND2–GLYCAN C1 angles between 110° and 130°.^12^

##### O-Glycosylation Site Geometric Validity

For *O*-linked glycoproteins, the SER CB–SER OG–GLYCAN C1 angle was calculated at detected *O*-glycosylation sites. For threonine residues, the THR CB–THR OG1–GLYCAN C1 angle was calculated. A structure passed the *O*-glycosylation site geometry check when all detected and scorable *O*-glycosylated sites had angles between 104° and 124°.^12^

##### Privateer N-Glycosite Scoring

Privateer glycosite torsion *Z*-scores^13^ were parsed from the Privateer output. The Privateer glycosylation score was defined as the fraction of evaluated samples that did not contain any glycosite torsion score below −1.0. All glycosite Z-score values below −1.0 were flagged as outliers.^13^

Some benchmarks (PDB Lectin, PDB Glycoprotein, Monosaccharide, and Oligosaccharide) evaluated both Top-1 and Oracle scores. For a given sample in these benchmarks, 5 predictions were generated. Top-1 refers to the prediction with the highest confidence score, while Oracle refers to the highest scoring prediction for each metric. For confidence score calculation, the following algorithm was used: confidence_score c = 0.8 * complex_plddt + 0.2 * iptm. Oracle metrics were computed across the evaluated samples for each target by selecting the best value for each metric independently: maximum for GlobalLDDT, LDDT-PLI, L-RMSD < 2.0, total glycan stereochemistry, dihedral < 0.1, sugar-RMSD<0.25, and the privateer glycosite validity score.

**Table S3.** Number of predicted samples used in each benchmark.

| **Benchmark Type** | **# of predicted samples** |
| --- | --- |
| PDB Lectin Benchmark | 5 |
| Untrained Lectin Benchmark | 1 |
| PDB Glycoprotein Benchmark | 5 |
| Untrained Glycoprotein Benchmark | 1 |
| Monosaccharide Benchmark | 5 |
| Oligosaccharide Benchmark | 5 |
| Untrained Oligosaccharide Benchmark | 1 |

